# Landscape of X chromosome inactivation across human tissues

**DOI:** 10.1101/073957

**Authors:** Taru Tukiainen, Alexandra-Chloé Villani, Angela Yen, Manuel A. Rivas, Jamie L. Marshall, Rahul Satija, Matt Aguirre, Laura Gauthier, Mark Fleharty, Andrew Kirby, Beryl B. Cummings, Stephane E. Castel, Konrad J. Karczewski, François Aguet, Andrea Byrnes, Consortium GTEx, Tuuli Lappalainen, Aviv Regev, Kristin G. Ardlie, Nir Hacohen, Daniel G. MacArthur

## Abstract

X chromosome inactivation (XCI) silences the transcription from one of the two X chromosomes in mammalian female cells to balance expression dosage between XX females and XY males. XCI is, however, characteristically incomplete in humans: up to one third of X-chromosomal genes are expressed from both the active and inactive X chromosomes (Xa and Xi, respectively) in female cells, with the degree of “escape” from inactivation varying between genes and individuals^1,2^ (Fig. 1). However, the extent to which XCI is shared between cells and tissues remains poorly characterized^3,4^, as does the degree to which incomplete XCI manifests as detectable sex differences in gene expression^5^ and phenotypic traits^6^. Here we report a systematic survey of XCI using a combination of over 5,500 transcriptomes from 449 individuals spanning 29 tissues, and 940 single-cell transcriptomes, integrated with genomic sequence data (Fig. 1). By combining information across these data types we show that XCI at the 683 X-chromosomal genes assessed is generally uniform across human tissues, but identify examples of heterogeneity between tissues, individuals and cells. We show that incomplete XCI affects at least 23% of X-chromosomal genes, identify seven new escape genes supported by multiple lines of evidence, and demonstrate that escape from XCI results in sex biases in gene expression, thus establishing incomplete XCI as a likely mechanism introducing phenotypic diversity^6,7^. Overall, this updated catalogue of XCI across human tissues informs our understanding of the extent and impact of the incompleteness in the maintenance of XCI.

---

XCI is an inherently random process, meaning that female tissues consist of two mixed cell populations, each with either the maternally or paternally inherited X chromosome marked for inactivation (Fig. 1). As such, assessments of XCI have often been confined to the use of artificial cell systems^1^, or samples presenting with skewed XCI^1,2^, i.e. preferential inactivation of one of the two X chromosomes, which is common in clonal cell lines but rare in karyotypically normal, primary human tissues^8^ (Supplementary Note and Extended Data Fig. 1). Others have used bias in DNA methylation^3,4,9^ or in gene expression^5,10^ between males and females as a proxy for XCI status. Here, we describe a systematic survey of the landscape of XCI, using a combination of three complementary approaches based on high-throughput mRNA sequencing (RNA-seq) (Fig. 1) that together allow an assessment of XCI from individual cells to population and across a diverse range of human tissues.

**Figure 1.**
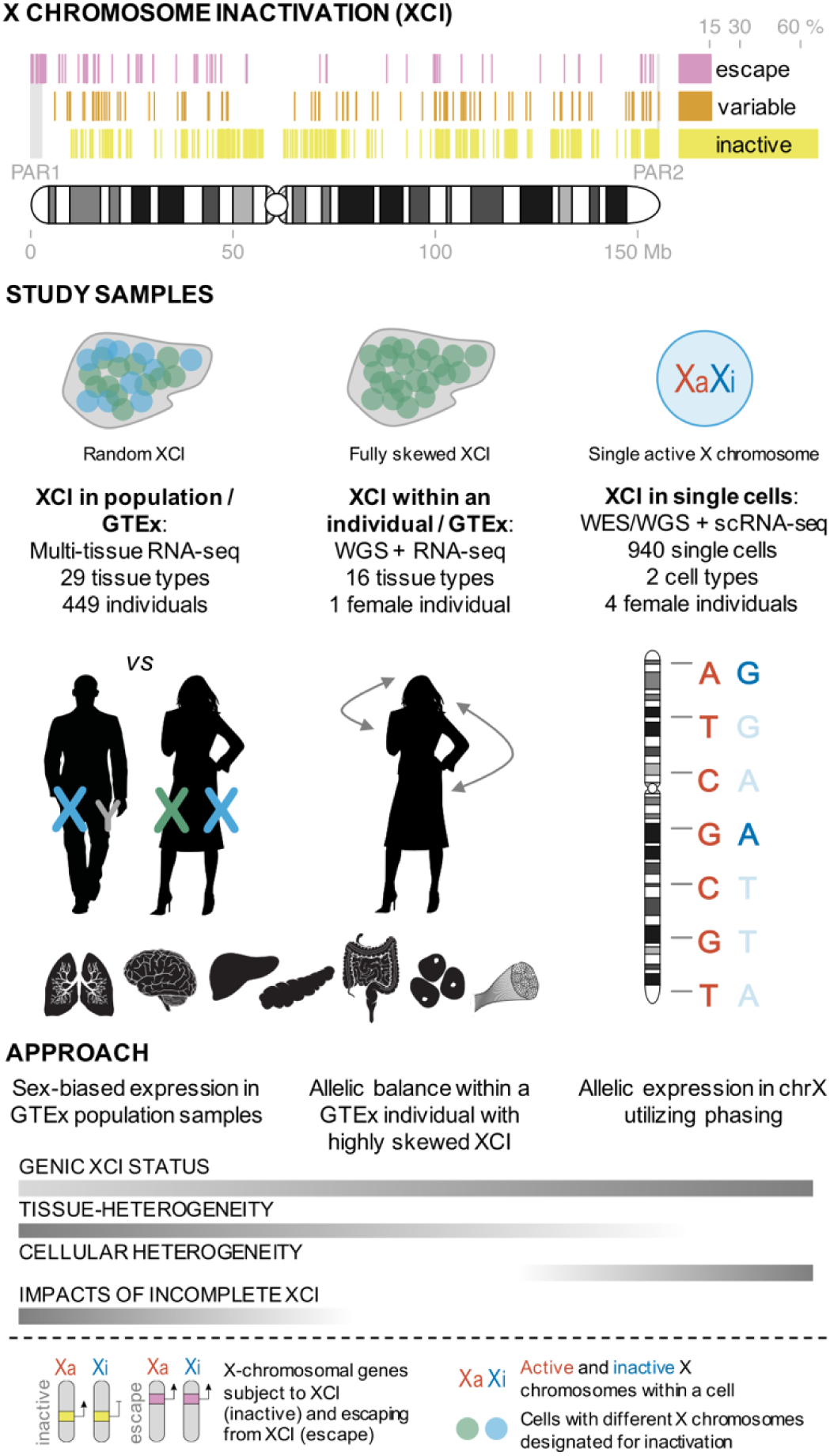
Schematic overview of the study. Previous expression-based surveys of XCI^1,2^ have established the incomplete and variable nature of XCI, showing that XCI is incomplete for up to 30% of X-chromosomal genes, but these studies have been limited in the tissue types and samples assessed. To investigate the landscape of XCI across human tissues, we combined three approaches: 1) sex-biases in expression using population-level GTEx data across 29 tissue types, 2) allelic expression in 16 tissue samples from a female GTEx donor with fully skewed XCI, and 3) validation using single cell RNA-seq by combining allelic expression and genotype phasing. WGS, whole genome sequencing; WES, whole exome sequencing; scRNA-seq, single-cell RNA-seq.

The limited accessibility of most human tissues, particularly in large sample sizes, means that no global investigation of the impact of incomplete XCI on X-chromosomal expression has been conducted in data sets spanning multiple tissue types. We thus used the Genotype Tissue Expression (GTEx) project^11^ data set, which includes high-coverage RNA-seq data from diverse human tissues, to investigate male-female differences in expression of 681 X-chromosomal protein-coding and long non-coding RNA (lncRNA) genes in 29 adult tissues (sample size per tissue 77-361; Extended Data Table 1, Methods) from the GTEx V6 data release, hypothesizing that escape from XCI should typically result in higher female expression of these genes. Previous work using expression microarrays^5^,^10^ has indicated that at least a subset of escape genes show a characteristic female bias in expression, but the higher sensitivity of deep RNA-seq and the more extensive set of profiled tissues in our analysis allows for the detection of subtler and tissue-specific phenomena.

To confirm that male-female expression difference serves as an accurate proxy for incomplete XCI, we assessed the enrichment of sex-biased expression in known XCI categories using 562 genes with a previous assignment of XCI status, defined as either escape (N=82), variable escape (N=89, i.e. genes where XCI status is variable between individuals) or inactive (N=391), combined from two previous expression-based surveys of XCI (Methods and Supplementary Table 1) (Fig. 1). As expected, sex-biased expression is more prominent among known escape genes compared to both inactive (two-sided Wilcoxon P=6.46×10^−11^) and variable escape genes (P=9.61×10^−11^) (Fig. 2b and Supplementary Table 2), with 74% of escape genes showing significant (false discovery rate (FDR) q-value < 0.01) male-female differences in at least one of the 29 tissues (Fig. 2a and Supplementary Table 3). In line with active transcription from two X-chromosomal copies in females, escape genes in the non-pseudoautosomal (nonPAR) region, i.e. in the X-specific region of the chromosome, are predominantly female-biased in expression across tissues (52 out of 67 assessed genes, binomial P=6.46×10^−6^). However, genes in the pseudoautosomal region in the tip of Xp, PAR1, are expressed more highly in males (14 out of 15 genes, binomial P=9.77×10^−6^) (Fig. 2a), suggesting that combined expression from Xa and Xi in females fails to reach the expression arising from both X and Y chromosomes in males, a result discussed in more detail below.

**Figure 2.**
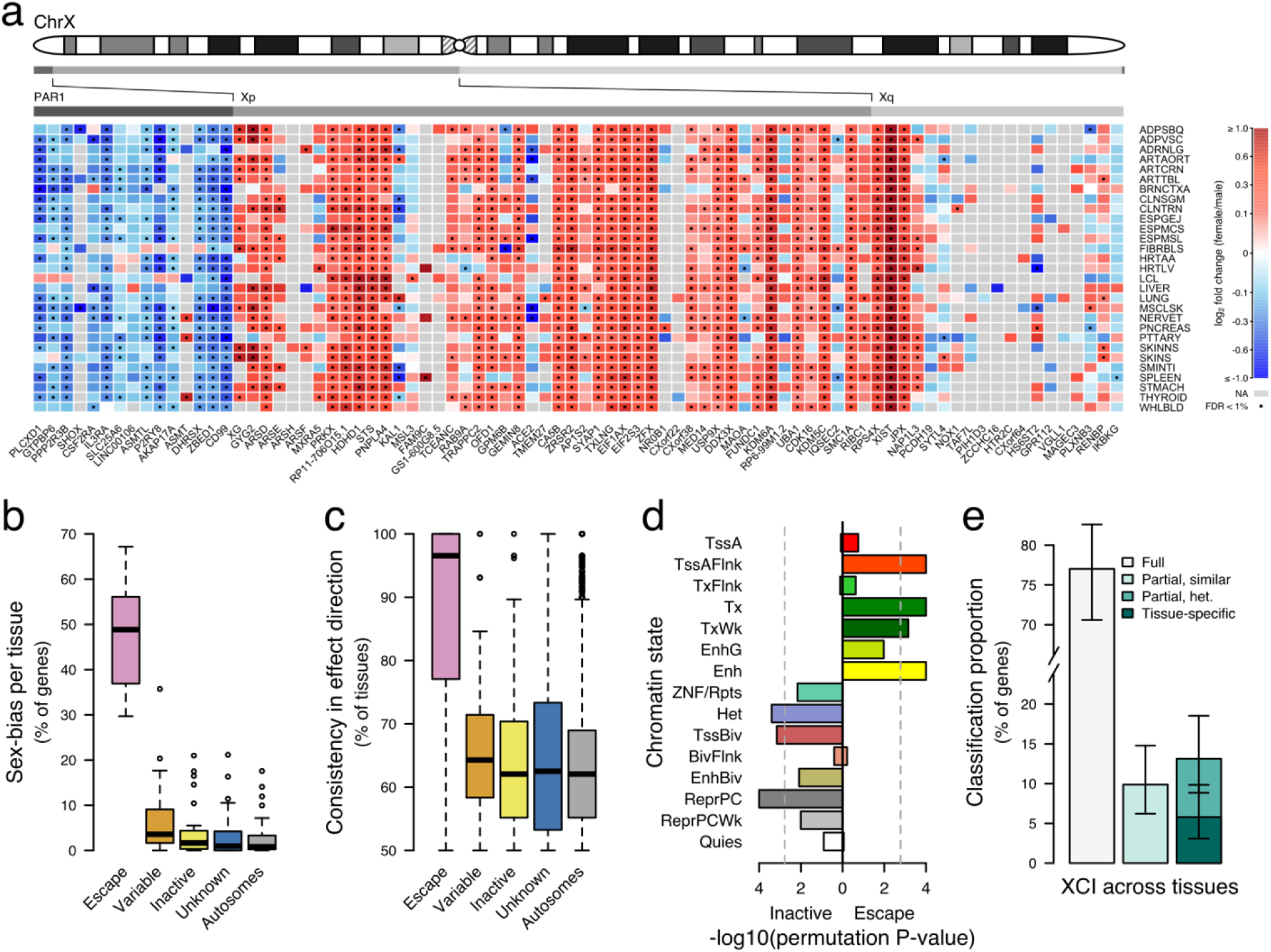
Assessment of tissue-sharing and population-level impacts of incomplete XCI. a) Heatmap representation of male-female expression differences in reported XCI-escaping genes (N=82) across 29 GTEx tissues. The color scale displays the direction of sex bias with red color indicating higher female expression. Genes that were too weakly expressed in the given tissue type to be assessed in the sex bias analysis are colored grey. Dots mark the observations where sex bias was significant at FDR<1%. b) Proportion of significantly biased (FDR<1%) genes in each tissue by reported genic XCI status. c) Proportion of tissues where the bias direction is shared by reported genic XCI status. Only genes expressed in at least five tissues are included. d) Chromatin state enrichment between reported escape and inactive genes in female samples from the Roadmap Epigenomics data. e) Classification of X-chromosomal genes (N=186) into full or incomplete and tissue-shared or heterogeneous XCI based on the analysis of chrX ASE patterns across tissues in a GTEx donor with fully skewed XCI. Error bars show 95% credible interval.

We find that sex bias of escape genes is often shared across human tissues, with these genes showing a higher number of tissues with significantly sex-biased expression than genes in other XCI categories (Fig. 2a and Supplementary Table 2). This result is not explained by a difference in the number of tissues in which escape and inactive genes are expressed (Supplementary Table 2), and there is marked consistency in the direction of sex bias across tissues (Fig. 2a,c and Supplementary Table 2). Together these observations point toward global and tight control of XCI. Previous reports have suggested that escape genes are unusual in their epigenomic landscape^12,13^; here we show that these genes are enriched in chromatin states related to active transcription (Fig. 2d) using the Roadmap Epigenomics Consortium^14^ data.

While consistent sex bias is remarkably specific to escape genes (Fig. 2b-c), a handful of genes show unexpected patterns. Nine genes without assigned XCI status, or previously annotated as inactive in some cell types, show more than 90% concordance in effect direction and significant sex bias (Extended Data Table 2). This includes *CHM*, for which we find independent evidence for incomplete XCI in single-cell RNA-seq (scRNA-seq; see below), and *RP11-706O15.3*, a lncRNA residing between X-Y homologous escape and variable escape genes *PRKX* and *NLGN4X* near PAR1 boundary, consistent with known clustering of escape genes^1,2^. Some escape genes show a more heterogeneous sex-biased pattern (Fig. 2a) suggesting either subtler or more tissue-specific escape, or alternatively hormonally regulated gene expression hampering the detection of the expected female bias (Supplementary Discussion). A cluster of such genes lies in the evolutionarily older region of the chromosome^15^, in Xq, where escape genes also show higher tissue-specificity and lower expression levels (Extended Data Table 3, Supplementary Discussion).

While sex bias is a broadly accurate proxy of XCI status, it provides only an indirect measure of the heterogeneity in inactivation across tissues. We serendipitously identified a GTEx female donor with an extremely unusual degree of skewing of XCI, resulting in the same copy of chrX being silenced in ~100% of cells across all tissues, yet without any X-chromosomal abnormality detected by whole-genome sequencing (WGS) (Methods, Supplementary Note and Extended Data Fig. 2). This provides an opportunity to validate the findings described above by leveraging allele-specific expression (ASE) in the 16 RNA-sequenced tissue samples available from this individual. We thus applied a statistical method designed for the comparison of patterns of ASE across tissues^16,17^ to the retrieved allelic counts (Supplementary Tables 4-6), which highlights the widespread incompleteness of XCI and the consistency of XCI across tissues, consistent with the preceding sex-bias analysis (Methods and Extended Data Table 4).

In this donor, approximately 23% of the 186 X-chromosomal genes expressed in at least two available tissues show evidence for expression from both alleles (Fig. 2e), aligning with previous estimates of the extent of escape from XCI^1,2^. For 43% of the biallelically expressed genes the expression arising from Xi is of similar magnitude between tissues (Fig. 2e), further supporting the observation of global and tight control of XCI from the sex-bias analysis, but the remainder display varying levels of Xi expression, suggesting a degree of tissue-dependence in XCI. In fact, this group of genes with more heterogeneous patterns includes a subset (5.8% of all genes) that appear biallelic in only one of the multiple tissues assayed. While analyses of mouse tissues have implicated several examples of tissue-specific escape^18^, previous methylation-based approaches have provided limited evidence for such a pattern in human tissues beyond neuronal samples^3,4,9^. As an example, in this sample, among the genes with the strongest probability for tissue-specific escape is *KAL1*, the causal gene for X-linked Kallman syndrome^19^; in line with strong female bias in lung expression in the GTEx sex bias analysis (Fig. 2a), in this individual *KAL1* shows biallelic expression exclusively in lung, thus likely representing the first validated example of a tissue-specific escape gene in humans. As a whole, the ASE-based predictions of XCI status in this sample align with previous assignments (Supplementary Table 7), yet the results also suggest six new escape genes (Extended Data Table 5), four of which act in a tissue-specific manner, e.g. *CLIC2* near the PAR2 boundary, illustrating the need for exploration of multiple tissue types to fully uncover the diversity in XCI.

While unprecedented in the breadth of tissue types analyzed, the above GTEx analyses share similar limitations to previous surveys^2,3,5^ in that they are confined to leveraging rare occurrences of XCI skewing or using indirect measures to assess patterns of XCI. The recent emergence of scRNA-seq methods^20–22^ presents an opportunity for studies of XCI at single cell resolution, removing the complication of the cellular heterogeneity in bulk tissue samples (Fig. 1). To establish the potential of scRNA-seq to directly profile XCI, we examined scRNA-seq data in combination with deep genotype sequences from 940 immune-related cells from four females: 198 cells from LCLs sampled from three females of African (Yoruba) ancestry, and 742 blood dendritic cells from a female of Asian ancestry (Villani et al., *co-submitted*) (Fig. 1, Extended Data Table 6). We utilized ASE to distinguish the expression coming from each of the two X-chromosomal haplotypes in a given cell. Studying allele-specific phenomena in single cells is complicated by widespread monoallelic expression, both biological and technological in origin^23^, which is present in autosomal as well as X-chromosomal genes^24–26^. To overcome this limitation, besides searching for X-chromosomal sites with biallelic expression (Extended Data Figure 3), we leveraged genotype phase information to detect sites where the expressed allele was discordant with the active X chromosome in that cell (Methods and Fig. 1).

We first confirmed that our single cell data replicates well-established XCI results. Across the four samples, 165 protein-coding and lncRNA genes (41-98 per sample) had informative heterozygous sites for analysis (Methods and Supplementary Table 4). While allelic dropout, which is extensive in scRNA-seq^21,23^, can lead to false negative calls in ASE (in our approach resulting in true escape genes being classified as inactivated), our XCI status estimates are robust to such errors and are overall consistent with previous observations: only 126 (76%) of the assayed genes were fully inactivated. The remaining genes showed incomplete XCI, i.e. Xi expression deviating significantly from baseline (Methods), in one or more samples (Fig. 3a-b, Supplementary Table 8), generally consistent with previous assignments of XCI escape status to these genes (Fig. 3a and Supplementary Table 8). For instance, single cell data reveal consistent expression from both X-chromosomal alleles for eleven genes in PAR1, in line with their known escape from XCI (e.g. *ZBED1* in Fig. 3c), and also replicate the known expression of *XIST* exclusively from the inactive X chromosome^27^ (Fig. 3d).

**Figure 3.**
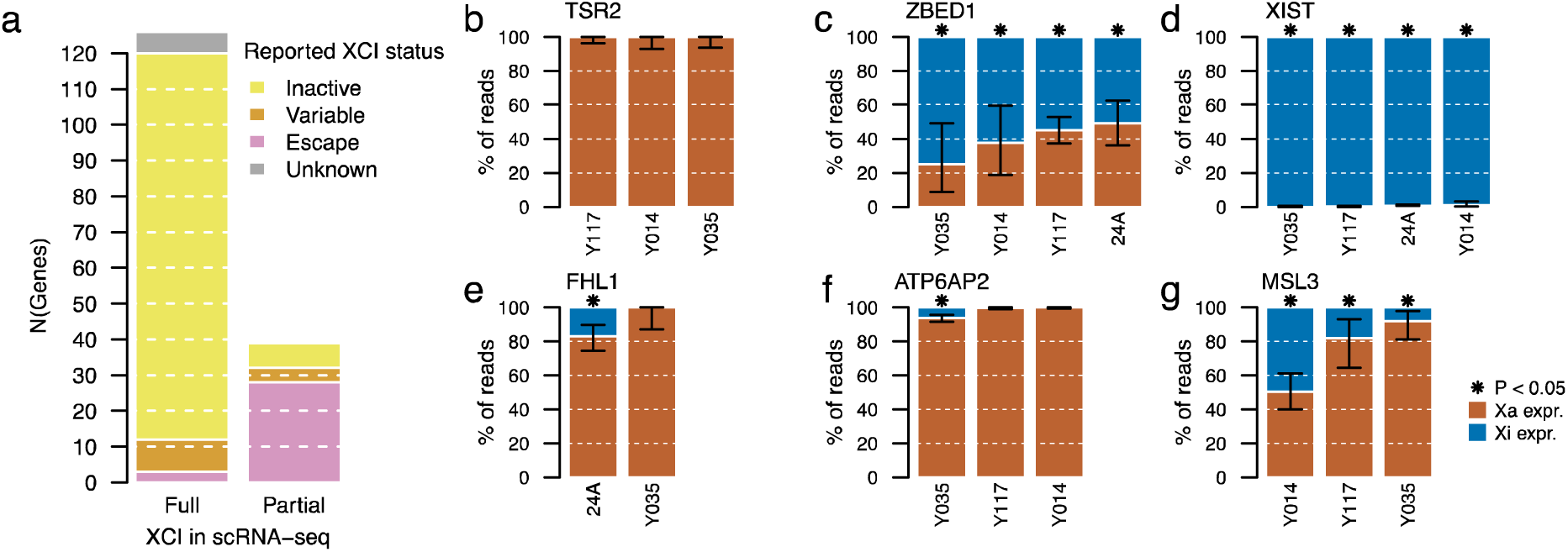
Analysis of XCI using scRNA-seq. a) Proportion of genes demonstrating full and partial XCI in the ASE analysis in single cell RNA-seq data from four individuals, and the corresponding concordance with previously reported XCI status. b-g) Examples of genes with different XCI patterns in scRNA-seq: previously reported inactive gene, which is inactive in all three scRNA-seq Yoruba samples (b), known escape gene in PAR1 (c), escape gene with known exclusive expression from Xi (d) new candidates for escape genes that demonstrate incomplete XCI in only a subset of samples (e and f), and a known escape gene that shows escape of varying degrees in the three samples (Pearson's Chi-squared test for equal proportions, P-value=3.80×10^−7^) (g). Asterisk above a bar indicates that the proportion of Xi expression, i.e. blue bar, in a given sample is significantly greater than the expected baseline (Binomial test P-value < 0.05; Methods). In (d) for 24A, i.e. the sample without parental genotype information, the allele expressed at *XIST* is assumed to be the Xi allele, in line with the exclusive Xi expression in the Yoruba samples confirmed using the information on parental haplotypes in each sample and the chromosome active in each cell.

Given the robustness of scRNA-seq in detecting known XCI-related phenomena, we next assessed whether our approach could further extend the spectrum of escape from XCI. For seven genes, the data from single cells conflicted with a previous assignment of the target gene's inactivation status (Supplementary Table 9), including *FHL1* (Fig. 3e), highlighted as a candidate escape gene also in the GTEx ASE analysis, and *ATP6AP2* (Fig. 3f), which displays predominantly female-biased expression across GTEx tissues. Both of these genes demonstrate significant Xi expression in only a subset of the scRNA-seq samples, a pattern consistent with variable escape^1,2,13^. Additionally, we show that between-individual variability exists not only in the presence but also in the degree of expression from Xi (e.g. *MSL3* in Fig. 3g). Further, highlighting the capacity of scRNA-seq to provide information beyond bulk RNA-seq, we identify examples where Xi expression varies considerably between the two X-chromosomal haplotypes within an individual (e.g. *ASMTL*; Supplementary Table 10), which suggests contribution from cis-acting regulatory variation as one of the determinants for the level of Xi expression^3^. Additionally, as a further layer of heterogeneity in Xi expression, we find a unique pattern at *TIMP1*, an established variable escape gene. Here expression arising from Xi across cells is not significant as a whole, yet exclusive to a subset of cells that express the gene biallelically (Extended Data Figure 3), thus suggesting cell-to-cell variability in propensity for escape, potentially attributable to variation in *TIMP1* promoter methylation^28^.

Combining ASE estimates from the scRNA-seq and GTEx analyses allows us to infer the magnitude of the incompleteness of XCI. We find that expression from Xi at escape genes rarely reaches levels equal to Xa, but on average remains at 33% of Xa expression, yet with wide variability along the chromosome (Supplementary Discussion and Extended Data Figure 4a) as shown in previous reports in specific tissue types^1,2^. While full escape would be required to balance the expression dosages between males and females in PAR1, Xi expression remains below Xa expression also in this region (mean Xi to Xa ratio ~0.80), suggesting partial spreading of XCI beyond X-chromosome-specific regions. We find no evidence for systematic up-or downregulation of Y chromosome expression in PAR1 (Extended Data Figure 4b, Methods and Supplementary Discussion) indicating that the consistent male-bias in PAR1 gene expression observed in the population-level analysis (Fig. 2a) is due to incomplete escape in PAR1 in females. Similarly, the partial Xi expression is responsible for six of the twelve assessed X-Y homologous genes in nonPAR^29^ (Methods), becoming male-biased when the expression arising from the Y chromosome counterpart is accounted for (Extended Data Figure 4c).

By combining diverse types and analyses of high-throughput RNA-seq data, we have systematically assessed the incompleteness and heterogeneity in XCI across 29 human tissues (Supplementary Table 11). We establish that scRNA-seq is suitable for surveys of XCI and, extending recent scRNA-seq-based analysis of the initiation of X-inactivation^30^, present the first steps towards understanding the cellular-level variability in the maintenance of XCI. Our phasing-based approach allows for the full use of low-coverage scRNA-seq, yet as any single individual and cell type is informative for restricted number of genes, larger data sets with more diverse cell types and conditions are required to fully profile XCI. We have thus utilized the multi-tissue GTEx data set to explore XCI in a larger number of X-chromosomal genes and to assess the tissue-heterogeneity and impacts of XCI on gene expression differences between the sexes.

These analyses show that incomplete XCI is largely shared between individuals and tissues, and extend previous surveys by pinpointing several examples of variability in the degree of XCI escape between cells, chromosomes, and tissues, as well as predicting at least seven new XCI-escaping genes supported by multiple analyses. In addition, our data demonstrate that escape from XCI results in sex-biased gene expression in at least 60 genes, and thus may well contribute to sex differences in health and disease (Supplementary Discussion). As a whole, these results highlight the between-female and male-female diversity introduced by incomplete XCI, the biological implications of which remain to be fully explored.

## Methods

### GTEx data

The GTEx project^11^ collected tissue samples from 554 postmortem donors (187 females, 357 males; age range 20-70), produced RNA sequencing from 8,555 tissue samples and generated genotyping data for up to 449 donors (GTEx Analysis V6 release). More details of methods can be found in Aguet et al. (Aguet et al., *co-submitted*). All GTEx data, including RNA, genome and exome sequencing data, used in the analyses described are available through dbGaP under accession phs000424.v6.p1, unless otherwise stated. Summary data and details on data production and processing are also available on the GTEx Portal (http://gtexportal.org).

### Single-cell samples

For the human dendritic cells samples profiled, the healthy donor (ID: 24A) was recruited from the Boston-based PhenoGenetic project, a resource of healthy subjects that are re-contactable by genotype^31^. The donor was a female Asian individual from China, of 25 years of age at the time of blood collection. She was a non-smoker, had normal BMI (height: 168.7cm; weight: 56.45kg; BMI: 19.8), and normal blood pressure (108/74). The donor had no family history of cancer, allergies, inflammatory disease, autoimmune disease, chronic metabolic disorders or infectious disorders. She provided written informed consent for the genetic research studies and molecular testing, as previously reported (Villani et al., *co-submitted*).

Daughters of three parent-child Yoruba trios from Ibadan, Nigeria, (i.e. YRI trios) collected as part of the International HapMap Project, were chosen for single-cell profiling both to maximize heterozygosity and due to availability of parental genotypes allowing for phasing. DNA and LCLs were ordered from the NHGRI Sample Repository for Human Genetic Research (Coriell Institute for Medical Research): LCLs from B-Lymphocyte for the three daughters (catalogue numbers: GM19240, GM19199, GM18518) and DNA extracted from LCLs for all members of the three trios (catalogue numbers: DNA: NA19240, NA19238, NA19239, NA19199, NA19197, NA19198, NA18518, NA18519, NA18520). These YRI samples are referred to by their family IDs: Y014, Y035 and Y117.

### Clinical muscle samples

To assess whether PAR1 genes are equally expressed from X and Y chromosomes we used a combination of skeletal muscle RNA sequencing and trio genotyping from eight male patients with muscular dystrophy, sequenced as part of an unrelated study. Patient cases with available muscle biopsies were referred from clinicians starting April 2013 through June 2016. All patients submitted for RNA-sequencing had previously available trio whole exome sequencing with one sample having additional trio whole genome sequencing. Muscle biopsies were shipped frozen from clinical centers via liquid nitrogen dry shipper and, where possible, frozen muscle was sectioned on a cryostat and stained with H&E to assess muscle quality as well as the presence of overt freeze-thaw artifact.

### Genotyping

The GTEx V6 release includes WGS data for 148 donors, including GTEX-UPIC. WGS libraries were sequenced on the Illumina HiSeqX or Illumina HiSeq2000. WGS data was processed through a Picard-based pipeline, using base quality score recalibration and local realignment at known indels. We used the BWA-MEM aligner for mapping reads to the human genome build 37 (hg19). SNPs and indels were jointly called across all 148 samples and additional reference genomes using GATK's HaplotypeCaller version 3.1. Default filters were applied to SNP and indel calls using the GATK's Variant Quality Score Recalibration (VQSR) approach. An additional hard filter InbreedingCoeff <= −0.3 was applied to remove sites that VQSR failed to filter.

WGS for one of the clinical muscle samples was performed on 500 ng to 1.5 ug of genomic DNA using a PCR-Free protocol that substantially increases the uniformity of genome coverage.

These libraries were sequenced on the Illumina HiSeq X10 with 151 bp paired-end reads and a target mean coverage of >30x, and processed similarly as above.

The Y117 trio (sample IDs NA19240 (daughter), NA19238 (mother), and NA19239 (father)) was whole-genome-sequenced as part of the 1000 Genomes project as described previously^32^. The VCF file containing the WGS-based genotypes for SNPs (YRI.trio.2010_09.genotypes.vcf.gz) was downloaded from the project's FTP site. To convert the genotype coordinates (in human genome build 36) in the original VCF to hg19 we used the liftover script (liftOverVCF.pl) and chain files provided as part of the GATK package.

WES was performed using Illumina's capture Exome (ICE) technology (Y035, Y014, 24A) or Agilent SureSelect Human All Exon Kit v2 exome capture (clinical muscle samples) with a mean target coverage of >80x. WES data was aligned with BWA, processed with Picard, and SNPs and indels were called jointly with other samples using GATK HaplotypeCaller package version 3.1 (24A, clinical muscle samples) or version 3.4 (Y035, Y014). Default filters were applied to SNP and indel calls using the GATK's Variant Quality Score Recalibration (VQSR) approach. A modified version of the Ensembl Variant Effect Predictor was used for variant annotation for all WES and WGS data. For trio WES or WGS data the genotypes of the proband were phased using the PhaseByTransmission tool of the GATK toolkit.

### Single cell data preparation and sequencing

[this section will be added in an upcoming revision]

### RNA-seq in GTEx

RNA sequencing was performed using a non-strand-specific RNA-seq protocol with poly-A selection of mRNA using the Illumina TruSeq protocol with sequence coverage goal of 50M 76 bp paired-end reads as described in detail previously^11^. The RNA-seq data, except for GTEX-UPIC, was aligned with Tophat version v1.4.1 to the UCSC human genome release version hg19 using the Gencode v19 annotations as the transcriptome reference. Gene level read counts and RPKMs were derived using the RNA-SeQC tool^33^ using the Gencode v19 transcriptome annotation. The transcript model was collapsed into gene model as described previously^11^. Read count and RPKM quantification include only uniquely mapped and properly paired reads contained within exon boundaries.

### RNA-seq alignment to personalized genomes

For the four single-cell samples and for GTEX-UPIC RNA-seq data was processed using a modification of the AlleleSeq pipeline^34,35^ to minimize reference allele bias in alignment. We first generated a diploid personal reference genome for each of the samples with the vcf2diploid tool^34^ including all heterozygous biallelic single nucleotide variants identified in WES or WGS either together with (YRI samples) or without (GTEX-UPIC, 24A) maternal and paternal genotype information. The RNA-seq reads were then aligned to both parental references using STAR^36^ version 2.4.1a in a per-sample 2-pass mode (GTEX-UPIC and YRI samples) or version 2.3.0e (24A) using hg19 as the reference. The alignments were combined by comparing the quality of alignment between the two references: for reads aligning uniquely to both references the alignment with the higher alignment score was chosen and reads aligning uniquely to only one reference were kept as such.

### RNA-seq of clinical muscle samples

Patient RNA samples derived from primary muscle were sequenced using the GTEx sequencing protocol^11^ with sequence coverage of 50M or 100M 76 bp paired-end reads. RNA-seq reads were aligned using STAR^36^ 2-Pass version v.2.4.2a using hg19 as the reference. Junctions were filtered after first pass alignment to exclude junctions with less than 5 uniquely mapped reads supporting the event and junctions found on the mitochondrial genome. The value for unique mapping quality was assigned to 60 and duplicate reads were marked with Picard MarkDuplicates (v.1.1099).

### Catalogue of X-inactivation status

In order to compare results from the ASE and GTEx analyses with previous observations on genic XCI status we collated findings from two earlier studies^1,2^ that represent systematic expression-based surveys into XCI. Each study catalogues hundreds of X-linked genes and together the data span two tissue types.

Carrel and Willard^1^ surveyed in total 624 X-chromosomal transcripts expressed in primary fibroblasts in nine cell hybrids each containing a different human Xi. In order to find the gene corresponding to each transcript, the primer sequences designed to test the expression of the transcripts in the original study were aligned to reference databases based on Gencode v19 transcriptome and hg19 using an in-house software (unpublished) (Supplementary Methods). In total 553 transcripts primer pairs were successfully matched to X-chromosomal Gencode v19 reference mapping together to 470 unique chrX genes (Supplementary Methods). These 470 genes were split into three XCI status categories (escape, variable, inactive) based on the level of Xi expression (i.e. the number of cell lines expressing the gene from Xi) resulting in 75 escape, 51 variable escape and 344 inactive genes.

Cotton et a^2^ surveyed XCI using allelic imbalance in clonal or near-clonal female LCL and fibroblast cell lines and provided XCI statuses for 508 genes (68 escape, 146 variable escape, 294 subject genes). The data was mapped to Gencode v19 using the reported gene names and their known aliases (Supplementary Methods), resulting in a list of XCI statuses for 506 X-chromosomal genes.

The results were combined by retaining the XCI status in the original study where possible (i.e. same status in both studies or gene unique to one study) and for genes where the reported XCI statuses were in conflict the following rules were applied: 1) A gene was considered “escape” if it was called escape in one study and variable in the other, 2) “variable escape” if classified as escape and inactive, and 3) “inactive” if classified as inactive in one study and variable escape in the other. The final combined list of XCI statuses consisted of 631 X-chromosomal genes including 99 escape, 101 variable escape and 431 inactive genes.

### Analysis of sex-biased expression

We conducted differential expression analyses to identify genes that are expressed at significantly different levels between male and female samples using 29 GTEx V6 tissues with RNA-seq and genotype data available from more than 70 individuals after excluding samples flagged in QC and sex-specific, outlier and highly correlated tissues^37^. We only included autosomal and X-chromosomal protein-coding or lncRNA genes in Gencode v19, and further removed all lowly-expressed genes. (Supplementary Methods and Extended Data Table 1).

Differential expression analysis between male and female samples was conducted using the voom-limma pipeline^38–40^ available as an R package through Bioconductor (https://bioconductor.org/packages/release/bioc/html/limma.html) using the gene-level read counts as input. We adjusted the analyses for age, three principal components inferred from genotype data using EIGENSTRAT^41^, sample ischemic time, surrogate variables^42, 43^ built using the sva R package^44^, and the cause of death classified into five categories based on the 4-point Hardy scale (Supplementary Methods).

To control the false discovery rate (FDR), we used the qvalue R package to obtain q-values applying the adjustment separately for the differential expression results from each tissue. The null hypothesis was rejected for tests with q-values below 0.01.

### XY homolog analysis

A list of Y-chromosomal genes with functional counterparts in the X chromosome, i.e. X-Y gene pairs, was obtained from Bellott et al^29^, which lists 19 ancestral Y chromosome genes that have been retained in the human Y chromosome. After excluding two of the genes (*MXRA5Y* and *OFD1Y*), which were annotated as pseudogenes by Bellot et al and further four genes (*SRY*, *RBMY*, *TSPY*, and *HSFY*) that according to Bellot et al have clearly diverged in function from their X-chromosomal homologs, the remaining 13 Y-chromosomal genes were matched with their X chromosome counterparts using gene pair annotations given in Bellot et al or by searching for known paralogs of the Y-chromosomal genes. To test for completeness of dosage compensation at the X-Y homologous genes, the sex-bias analysis in GTEx data was repeated replacing the expression of the X-chromosomal counterpart with the combined expression of the X and Y homologs.

### Chromatin state analysis

To study the relationship between chromatin states and XCI, we used chromatin state calls from the Roadmap Epigenomics Consortium^14^. Specifically, we used the chromatin state annotations from the core 15-state model, publicly available at http://egg2.wustl.edu/roadmap/web_portal/chr_state_learning.html#core_15state. We followed our previously published method^45^ to calculate the covariate-corrected percentage of each gene body assigned to each chromatin state. After pre-processing, we filtered down to the 399 inactive and 86 escape genes on the X chromosome, and down to 38 female epigenomes.

To compare the chromatin state profiles of the escape and inactive genes in female samples, we used the one-sided Wilcoxon rank sum test. Specifically, for each chromatin state, we averaged the chromatin state coverage across the 38 female samples for each gene, and compared that average chromatin state coverage for all 86 escape genes to the average chromatin state coverage for all 399 inactive genes. We performed both one-sided tests, to test for enrichment in escape genes, as well as enrichment in inactive genes.

Next, we performed simulations to account for possible chromatin state biases, such as the fact that the escape and inactive genes are all from the X chromosome. Specifically, we generated 10,000 randomized simulations where we randomly shuffled the “escape” or “inactive” labels on the combined set of 485 genes, while retaining the sizes of each gene set. For each of these simulated “escape” and “inactive” gene sets, we calculated both one-sided Wilcoxon rank sum test p-values as described above, and then, we calculated a permutation “p-value” for the real gene sets based on these 10,000 random simulations (Supplementary Methods). Finally, we used Bonferroni multiple hypothesis correction for our significance thresholds to correct for our 30 tests, one for each of 15 chromatin states, and both possible test directions.

### Allele-specific expression

For ASE analysis the allele counts for biallelic heterozygous variants were retrieved from RNA-seq data using GATK ASEReadCounter (v.3.6)^35^. Heterozygous variants that passed VQSR filtering were first extracted for each sample from WES or WGS VCFs using GATK SelectVariants. The analysis was restricted to biallelic SNPs due to known issues in mapping bias in RNA-seq against indels^35^. Sample-specific VCFs and RNA-seq BAMs were inputted to ASEReadCounter requiring minimum base quality of 13 in the RNA-seq data (scRNA-seq samples, GTEX-UPIC) or requiring coverage in the RNA-seq data of each variant to be at least 10 reads, with a minimum base quality of 10 and counting only reads with unique mapping quality (MQ = 60) (clinical muscle samples).

For downstream processing of the scRNA-seq and GTEX-UPIC ASE data, we applied further filters to the data to focus on exonic variation only and to conservatively remove potentially spurious sites (Supplementary Methods), e.g. we removed sites with non-unique mappability, and further after an initial analyses of the ASE data subjected 22 of the X-chromosomal ASE sites to manual investigation. For GTEX-UPIC we limited the X-chromosomal ASE data in case of multiple ASE sites to only one site per gene, by selecting the site with coverage >7 reads in the largest number of tissues, to have equal representation from each gene for downstream analyses.

### Assessing ASE across tissues

For GTEX-UPIC sample for which we had ASE data from up to 16 tissues per each ASE site, we applied the two-sided Hierarchical Grouped Tissue Model (GTM*) implemented in MAMBA 1.0.0^16,17^ to ASE data. The Gibbs sampler was run for 200 iterations with a burn-in of 50 iterations.

GTM* is a Bayesian hierarchical model that borrows information across tissues and across variants, and provides parameter estimates that are useful for interpreting global properties of variants. It classifies the sites into ASE states according to their tissue-wide ASE profiles and provides an estimate of the proportion of variants in each of the five different ASE states (strong ASE across all tissues (SNGASE), moderate ASE across all tissues (MODASE), no ASE across all tissues (NOASE), and heterogeneous ASE across tissues (HET1 and HET0)).

To summarize the GTM* output in the context of XCI, we considered SNGASE to reflect full XCI, MODASE and NOASE together to represent partial XCI with similar effects across tissues, and HET1 and HET0 to reflect partial yet heterogeneous patterns of XCI across tissues. In order to combine estimates from two ASE states, we summed the estimated proportions in each class, and subsequently calculated the 95% confidence intervals for each remaining ASE state using Jeffreys prior.

### Biallelic expression in single cells

Biallelic expression in individual cells in the X chromosome was assessed only at ASE sites covered by the minimum of eight reads. We considered sites biallelically expressed when 1) allelic expression > 0.05, and 2) one-sided binomial test indicated allelic expression to be at least nominally significantly greater than 0.025. Only genes with at least two observations of biallelic expression across all cells within a sample were counted as biallelic.

### Phasing scRNA-seq data

We assigned each cell to either of two cell populations distinguished by the parental X-chromosome designated for inactivation utilizing genotype phasing. For the YRI samples, where parental genotype data was available, the assignment to the two parental cell populations was unambiguous for all cells where X-chromosomal sites outside PAR1 or frequently biallelic sites were expressed. For 24A no parental genotype data was available, and hence we utilized the correlation structure of the expressed X-chromosomal alleles across the 948 cells to infer the two parental haplotypes utilizing the fact that in individual cells the expressed alleles at the chrX sites subject to full inactivation (i.e. the majority chrX ASE sites), are from the X chromosome active in each cell (Supplementary Methods). For this calculation we excluded all PAR1 sites and all additional sites that were frequently biallelic, to minimize the contribution of escape genes to the phase estimation. After assigning each informative cell to either of the parental cell populations, the reference and alternate allele reads for each ASE site were mapped to active and inactive allele reads within each sample using the actual or inferred parental haplotypes. The data was first combined per variant by taking the sum of active and inactive counts separately across cells, and further similarly combined per gene, if multiple SNPs per gene were available.

### Determining XCI status from scRNA-seq ASE

Before calling XCI status using the Xa and Xi read counts, we excluded all sites without fewer than five cells contributing ASE data at each gene and also all sites with coverage lower than eight reads across cells within each sample. To determine whether the observed Xi expression is significantly different from zero, hence indicative of incomplete XCI at the site / gene, we required two criteria to be fulfilled: 1) The Xi to total expression ratio above 0.05 for YRI samples and 0.075 for 24A (the higher threshold for 24A was selected to account for the larger uncertainty in assigning cells to the parental cell populations), and 2) the Xi to total expression ratio to be significantly (P<0.05) greater than hypothesized upper bound for error (determined as 0.025 for YRI samples and 0.05 for 24A) in a one tailed binomial test. Genes where at least one of the samples showed significant Xi expression were considered partially inactivated, while the remaining were classified as subject to full XCI.

### ChrX and chrY expression in PAR1

Using the parental origin of each allele reference and alternate allele read counts at PAR1 ASE sites were assigned to X and Y chromosomes (i.e. maternally and paternally inherited alleles, respectively). For each sample, the PAR1 ASE data was summarized by gene by taking the sum of X and Y chromosome reads across all informative ASE sites within each gene. Significance of deviation from equal expression was assessed using a two-sided binomial test.

### Manual curation of heterozygous variants from ASE analyses

Twenty-two heterozygous variants assessed in chrX ASE analysis were subjected to manual curation due to providing results in the XCI analysis that were in conflict with previous assignment of the underlying gene to be subject to full XCI. For each sample, BWA-aligned germline bams were viewed in IGV using either WGS or WES data. The presence of a number of characteristics called into question the confidence of the variant read alignments and thus the variant itself (Supplementary Methods). Allele balance that deviated significantly from 50:50 was considered suspect and often coincided with the existence of homology between the reference sequence in the region surrounding the variant and another area of the genome, as ascertained using the UCSC browser self-chain track and/or BLAT alignment of variant reads from within IGV. Other sequence-based annotations added to the VCF by HaplotypeCaller were also evaluated in the interests of examining other signatures of ambiguous mapping. The phasing of nearby variants was also considered. If phased variants occurred in the DNA sequencing data that were not assessed in the ASE analysis, those variants were considered suspect.

## Acknowledgements

We thank J. Maller, F. Zhao, and M. Lek for technical assistance and P. J. Siponen for assistance with figure design.

T.T. was supported by the Academy of Finland (285725), Finnish Cultural Foundation, Orion-Farmos Research Foundation, and Emil Aaltonen Foundation. K.J.K. is supported by NIGMS

Fellowship (F32GM115208). This work was supported by NIH grants U54DK105566, R01MH101820 and R01GM104371 to D.G.M.

The Genotype-Tissue Expression (GTEx) Project was supported by the Common Fund of the Office of the Director of the National Institutes of Health. Additional funds were provided by the NCI, NHGRI, NHLBI, NIDA, NIMH, and NINDS. Donors were enrolled at Biospecimen Source Sites funded by NCI\SAIC-Frederick, Inc. (SAIC-F) subcontracts to the National Disease Research Interchange (10XS170), Roswell Park Cancer Institute (10XS171), and Science Care, Inc. (X10S172). The Laboratory, Data Analysis, and Coordinating Center (LDACC) was funded through a contract (HHSN268201000029C) to The Broad Institute, Inc; this grant also provided funding to D.G.M. and T.T. Biorepository operations were funded through an SAIC-F subcontract to Van Andel Institute (10ST1035). Additional data repository and project management were provided by SAIC-F (HHSN261200800001E). The Brain Bank was supported by a supplements to University of Miami grants DA006227 & DA033684 and to contract N01MH000028. Statistical Methods development grants were made to the University of Geneva (MH090941 & MH101814), the University of Chicago (MH090951, MH090937, MH101820, MH101825), the University of North Carolina-Chapel Hill (MH090936 & MH101819), Harvard University (MH090948), Stanford University (MH101782), Washington University St Louis (MH101810), and the University of Pennsylvania (MH101822).

## Author Contributions

T.T. and D.G.M. designed the study. A.C.V. designed and conducted the scRNA-seq experiments. T.T., A.Y., M.A.R., M.A., L.G., M.F., and B.B.C. analyzed the data. J.L.M., R.S., S.E.C., A.K., K.J.K., F.A., A.B., T.L., A.R., K.G.A., N.H., and D.G.M. provided tools and reagents. T.T. and D.G.M wrote the manuscript with input from other authors.

